# Comparison of real-time PCR and droplet digital PCR for the detection of *Xylella fastidiosa* in plants

**DOI:** 10.1101/582288

**Authors:** Enora Dupas, Bruno Legendre, Valérie Olivier, Françoise Poliakoff, Charles Manceau, Amandine Cunty

## Abstract

*Xylella fastidiosa* (*Xf*) is a quarantine plant pathogen bacterium originating from the Americas and that has emerged in Europe in 2013. *Xf* can be detected directly on plant macerate using molecular methods such as real-time PCR, which is a sensitive technique. However, some plants may contain components that can act as PCR reaction inhibitors, which can lead to false negative results or an underestimation of the bacterial concentration present in the analyzed plant sample. Droplet digital PCR (ddPCR) is an innovative tool based on the partitioning of the PCR reagents and the DNA sample into thousands of droplets, allowing the quantification of the absolute number of target DNA molecules present in a reaction mixture, or an increase of the detection sensitivity. In this study, a real-time PCR protocol, already used for *Xf* detection in the framework of official surveys in the European Union, was transferred and optimized for *Xf* detection using ddPCR. This new assay was evaluated and compared to the initial real-time PCR on five plant matrices artificially inoculated and on naturally infected plants. In our conditions, this new ddPCR enabled the detection of *Xf* on all artificially inoculated plant macerates with a similar limit of detection, or a slight benefit for *Quercus ilex*. Moreover, ddPCR improved diagnostic sensitivity as it enabled detection of *Xf* in samples of *Polygala myrtifolia* or *Q. ilex* that were categorized as negative or close to the limit of detection using the real-time PCR. Here, we report for the first time a ddPCR assay for the detection of the bacterium *Xf*.

## 1 Introduction

*Xylella fastidiosa* (*Xf*) is a plant pathogenic bacterium known worldwide. Located in the xylem vessels of plants, its natural way of transmission is sap-feeding insect vectors (Almeida and Nunney, 2015). To date, five different subspecies (subsp.) have been described: subsp. *fastidiosa*, subsp. *morus*, subsp. *multiplex*, subsp. *pauca*, and subsp. *sandyi* (Nunney *et al*., 2014; Schaad *et al*., 2004; Schuenzel *et al*., 2005). In Europe, *Xf* was first detected in Italy, in the Apulia area in 2013, where the subsp. *pauca* was identified on olive trees (Saponari *et al*., 2013). Then, in France in 2015 (Corsica and in French Riviera region), the subsp. *multiplex* was reported firstly on *Polygala myrtifolia* and then on a large range of ornamental or wild plants (Denancé *et al*., 2017). More recently in 2016, the subsp. *fastidiosa, multiplex* and *pauca* were identified in the Balearic Islands (Spain), on olive trees, grapevines and sweet cherry trees; and in 2017, the presence of the subsp. *multiplex* was also identified in Spain near Alicante, on almond trees (Landa, 2017). Currently, 563 plant species distributed in 82 botanical families are reported to be hosts of *Xf*, and this list includes plants of major socio-economic interest such as olive trees, citrus or grapevine (EFSA, 2018). Since 2017, *Xf* has been classified in Annex I/A2 of Council Directive 2000/29/EC revised in 2017, and in the A2 list of the EPPO as a quarantine pathogen present on the EU territory and requiring mandatory control (C/2017/4883, 2017; EPPO, 2018a).

As isolation and cultivation of *Xf* is fastidious, detection and identification tests are applied directly on plant extracts (Denancé *et al*., 2017). Nowadays, different molecular tools targeting specific DNA regions are available to detect the bacterium at the species level or to specifically detect one of the subspecies. Conventional PCRs such as Minsavage *et al*. (2014) have been developed, but they are less sensitive than Real-Time PCR (Baldi and La Porta, 2017). Among the real-time PCR techniques developed, the method designed by Harper *et al*. (2010) was identified as one of the most suitable methods for *Xf* detection. It allow to detect all the *Xf* subspecies, its limit of detection determined on different plant species is low, it is sensitive, and no cross-reactions with other bacterial species have been reported (Francis *et al*., 2006; Harper *et al*., 2010; Li *et al*., 2013; Modesti *et al*., 2017; Ouyang *et al*., 2013).

Even though real-time PCR is very sensitive in most cases, low bacterial contamination levels of plants and the presence of PCR inhibitors can lead to false negative results, and the underestimation of positive samples for some plant species (Modesti *et al*., 2017; Schrader *et al*., 2012). These PCR inhibitors include polyphenols, polysaccharides, pectin and xylan (Harper *et al*., 2010; Schrader *et al*., 2012; Wei *et al*., 2008). Studies have revealed that the improvement of DNA extraction methods, or the addition of bovine serum albumin (BSA) during the PCR assay, may reduce the impact of PCR inhibitors (Harper *et al*., 2010; Schrader *et al*., 2012). For some plants, such as *Nerium oleander, Prunus dulcis* and *Vitis vinifera*, a tenfold dilution prevented PCR inhibition and led to successful detection of *Xf* (Francis *et al*., 2006; Minsavage *et al*., 1994; Modesti *et al*., 2017). However, DNA dilution cannot be applied to every sample, due to low *Xf* concentrations in some infected plants. It can therefore be rather difficult to find a universal method, because of the wide range of *Xf* host plants. Moreover, even though real-time PCR produces quantitative data when using a calibration curve, the results are often only interpreted qualitatively for *Xf* detection (Cruaud *et al*., 2018; Modesti *et al*., 2017).

Digital PCR was set up in 1999 by Vogelstein and Kinzler and named later by Morley in 2014. By compartmentalizing the PCR reaction into thousands of droplets, ddPCR offers the promises of absolute quantification without the need for calibration (Hindson *et al*., 2011; Huggett *et al*., 2013; Morley, 2014; Voegel and Nelson, 2018; Vogelstein and Kinzler, 1999). At first, ddPCR was designed to identify rare mutations in a small number of cells (Vogelstein and Kinzler, 1999). Already used for medical purposes (Bharuthram *et al*., 2014; Cao *et al*., 2015; Hindson *et al*., 2011; Nixon *et al*., 2014; Ramírez *et al*., 2019), this method was transferred as a detection and quantification tool to other fields, such as environmental sciences (Doi *et al*., 2015; Hoshino and Inagaki, 2012), food safety control (Bian *et al*., 2015; Wang *et al*., 2018), GMO detection (Košir *et al*., 2017; Morisset *et al*., 2013) or the agronomic field (Dreo *et al*., 2014; Maheshwari *et al*., 2017; Rački *et al*., 2014; Voegel and Nelson, 2018; Zhao *et al*., 2016). The transfer of real-time PCR assays to ddPCR assays has already provided successful results for the detection and the quantification of plant pathogenic bacteria (Dreo *et al*., 2014; Maheshwari *et al*., 2017; Zhao *et al*., 2016). For example, ddPCR was more efficient than real-time PCR to detect low concentrations of *Ralstonia solanacearum* in potatoes (Dreo *et al*., 2014). It also increased the detection threshold of other pathogens such as *Xanthomonas citri* subsp. *citri* in citrus, Pepper mild mottle virus in plants, soil and water, or of GMOs in maize seed powder (Morisset *et al*., 2013; Rački *et al*., 2014; Zhao *et al*., 2016). ddPCR was reported to be up to 1,000 fold more sensitive than conventional PCR developed to detect adenovirus in live attenuated vaccines (Dong *et al*., 2018). It allows the detection and quantification of pathogen abundance, such as *Agrobacterium vitis* in grapevines, for which previous methods lacked sensitivity (Voegel and Nelson, 2018). An additional advantage of ddPCR is its tolerance to PCR inhibitors present in plants, soil, water or food (Cao *et al*., 2015; Maheshwari *et al*., 2017; Morisset *et al*., 2013; Rački *et al*., 2014; Zhao *et al*., 2016). ddPCR presents many advantages that could make it an alternative for *Xf* detection.

The aim of this study was to transfer the real-time PCR developed by Harper *et al*. (2010) into a ddPCR assay, in order to improve the detection of *Xf* at low concentrations in plant matrices rich in PCR inhibitors. ddPCR was compared to real-time PCR using five artificially inoculated plant matrices and naturally infected plants sampled in France. The plant species used as matrices were selected for their level of PCR inhibitors reported by the Plant Health Laboratory (PHL) of the French Agency for Food, Environmental and Occupational Health & Safety (ANSES) following the analysis of thousands of different plant samples collected since 2015 in the context of the national *Xf* survey in France (personal communication, Bruno Legendre). *Polygala myrtifolia* was selected as a matrix containing a low concentration of inhibitors, *Lavandula angustifolia, Olea europaea, Quercus ilex* and *Rosmarinus officinalis* were selected as matrices containing high concentrations of inhibitors. Experimental assays were set up following the standard PM7/98 and the digital MIQE guidelines. The following performance criteria were evaluated: analytical sensitivity, repeatability, and diagnostic specificity (EPPO, 2014; Huggett *et al*., 2013).

## 2 Materials and methods

### 2.1 Bacterial strains

Bacterial strains *Xf* subsp. *multiplex* CFBP 8416, isolated in France (Corsica) in 2015 from symptomatic *P. myrtifolia* (Denancé *et al*., 2017) and *Xf* subsp. *fastidiosa* CFBP 7970, isolated in the United States (Florida) from *Vitis vinifera* were cultivated on modified PWG media at 28°C for two weeks (EPPO, 2018b). Bacterial suspensions of pure culture of *Xf* were prepared in sterile demineralized water and suspensions titer was estimated by immunofluorescence (IF) (EPPO, 2009, 2018b). The antiserum, used to count *Xf*, was especially produced in collaboration with the UR1268 BIA - Team Allergy of the French National Institute for Agricultural Research (INRA) of Angers-Nantes. It resulted from the inoculation of rabbits with nine strains of *Xf* chosen to be as diverse as possible in terms of subspecies, geographical location, and host plant species. The initial concentration of the CFBP 8416 bacterial suspension was estimated by IF at 1.84×10^9^ bacteria/mL (b/mL). This suspension was used to spike all the artificially inoculated samples in this study. The bacterial suspension of the strain CFBP 7970 was calibrated at 1×10^7^ b/mL and used as a positive control for the PCR reactions.

### 2.2 Plant materials

Healthy plant materials used for spiking assays were collected in 2018. *L. angustifolia, O. europaea, Q. ilex, R. officinalis* were sourced from Maine-et-Loire, a French department known to be *Xf* free. *P. myrtifolia* was produced in a nursery in Brittany (*Xf* free) and had a European phytosanitary passport certifying its healthy status. Moreover, no symptoms were recorded on these five plants. In this study, the healthy status of the five matrices was first checked using the real-time PCR assay Harper *et al*. (2010).

Naturally infected samples of *Calicotome* sp. (one sample), *L. angustifolia* (four samples), *P. myrtifolia* (13 samples), and *Q. ilex* (4 samples) were collected in the context of the national survey, between 2016 and 2018 in Corsica and in the PACA region of France. These 22 samples were already found to have a positive status or to be at the limit of detection by the PHL, using the real-time PCR developed by Harper *et al*. (2010).

### 2.3 Plant spiking

The artificially inoculated plant samples were prepared by mixing 1 g of healthy plant petiole in 4.5 mL of sterile distilled water and spiked with 0.5 mL of a known concentration of bacterial suspension. Each matrix was spiked in order to reach a range dilution of 1×10^5^ b/mL; 5×10^4^ b/mL; 1×10^4^ b/mL; 5×10^3^ b/mL; 1×10^3^ b/mL; 5×10^2^ b/mL; and 1×10^2^ b/mL. The negative template control (NTC) was obtained by mixing 1 g of healthy plant petiole with 5 mL of sterile distilled water.

### 2.4 DNA extraction

The bacterial strain suspension used as a positive control for all the PCRs was inactivated by thermal lysis. A volume of 1 mL of bacterial suspension was heated at 100°C for 5 min and then frozen at −20°C for at least 15 min.

The macerates of all the spiked plants and naturally infected samples were crushed, prior to DNA extraction. DNA extractions and purifications were carried out using the QuickPick™ SML Plant DNA Kit (Bio-Nobile, Turku, Finland). Extraction, washing, and elution of the DNA were automated using KingFisher™ mL (Thermo Scientific). DNA extracts were kept at 4°C for a week, or stored at −20°C for a longer period.

### 2.5 Real-time PCR Harper et al., 2010

The real-time PCR assays were performed on the thermal cycler CFX96 real-time System C1000 Touch (Bio-Rad), using 96-well plates (Hard-Shell^®^ 96-Well PCR Plates, #hsp9601, Bio-Rad). The following thermal cycling program used was: 50°C for 2 min, 94°C for 10 min, then 40 cycles of two step of 94°C for 10 s and 62°C for 40 s. The reaction mix was prepared in a final volume of 20 µL containing: 1x TaqMan Fast Universal Master Mix (Applied Biosystems), 300 nM of each *Xf* forward and reverse primers (*XF*-F and *XF*-R, respectively), 100 nM of 6’FAM/BHQ-1 labeled probe (*XF*-P), 300 µg/µL of BSA, and 2 µL of DNA sample. For the artificially contaminated plant material, each sample was amplified in triplicate and on three independent PCR runs to obtain nine Ct values per sample. For the naturally infected plant material, each sample was amplified in duplicate on the same PCR run.

The data acquisitions and data analyses were performed using Bio-Rad CFX Manager, v 3.0. The determination of Ct values was done using the regression mode. A Ct higher than 38 was considered to be a negative result, according to the cut-off indicated by (Harper *et al*., 2010). For all the following analyses, the limit of detection was fixed at 100%, meaning the lowest concentration at which all replicates gave a positive signal.

### 2.6 Optimization and evaluation of the ddPCR assay

Two thermal gradients were tested to determine the optimal hybridization temperature ranging from 54.6 to 64.6°C, and from 57 to 62°C. Thermal gradients were applied on samples of *P. myrtifolia*, spiked with suspensions of *Xf* ranging from 1×10^5^ b/mL to 1×10^3^ b/mL. BSA was tested at the concentration determined by Harper *et al*. (2010) (300 µg/µL) on a sample of *P. myrtifolia* spiked with *Xf* at a concentration of 1×10^5^ b/mL. In order to optimize the assay for the five matrices spiked at 1×10^5^ b/mL, four different DNA volumes added to the reaction mix were tested: 2 µL; 4 µL; 6 µL and 8 µL. Using the optimized protocol, tenth dilutions of *L. angustifolia* and *R. officinalis* spiked with 1×10^3^ b/mL were tested. All the experiments conducted to optimize the ddPCR assay were amplified in triplicate.

### 2.7 Optimized ddPCR assay

The optimized ddPCR reaction mix conditions retained were a final reaction volume of 20 µL containing: 1x ddPCR™ Supermix for Probes (No dUTP) (Bio-Rad), 900 nM *Xf* forward and reverse primers (*XF*-F and *XF*-R), 250 nM 6’FAM/BHQ-1 labeled probe (*XF*-P), and 8 µL of DNA sample. Droplets were generated with the QX200™ Droplet Digital™ System (Bio-Rad) in a cartridge containing 20 µL of the reaction mix and 70 µL of Droplet Generation Oil for Probes (ddPCR™ 96-Well Plates #12001925, Bio-Rad). The entire emulsion volume was transferred from the cartridge to a 96-well PCR plate (Bio-Rad) and the PCRs were performed on the thermal cycler CFX96 real-time System C1000 Touch (Bio-Rad). Optimal thermocycling conditions retained were: DNA polymerase activation of 95°C for 10 min, then 40 cycles of two-steps of 94°C for 30 s for denaturation and 59°C for 60 s for hybridization and elongation, followed by a final step at 98°C for 10 min for droplet stabilization. According to Bio-Rad recommendations, a temperature ramp of 2°C/s was fixed on all PCR steps and the lead was heated at 105°C. After amplification, the PCR plate was directly transferred to the droplet reader QX200™ Droplet Digital™ System (Bio-Rad). QuantaSoft 1.7.4.0917 software was used for data acquisition and data analysis. The entire concentration range of spiked matrices was first amplified in triplicate on the same run. Then, for each matrix, samples with concentrations equal to and below the limit of detection identified by real-time PCR were amplified in triplicate on two other independent runs, in order to ultimately obtain nine results for these samples. Finally, the amplifications of the naturally infected samples were performed in one replicate in one run.

### 2.8 ddPCR analysis

Data were analyzed directly with QuantaSoft™ Analysis Pro software. For each plate and each matrix, a threshold was manually set up just above the amplitude value of the cloud corresponding to the negative droplets, also considered as the background, and according to the results of the corresponding NTC (Lievens *et al*., 2016). This threshold enabled the differentiation of droplets by categorizing them as positive (high level of fluorescence) or negative (low and constant level of fluorescence). The PCR reactions with fewer than 10,000 droplets generated were excluded from the analysis, and a result was considered positive if at least two positive droplets were detected. The software provided the results in target copies by reaction using the following formula:

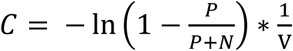

Where C corresponded to the concentration in target DNA copies/well (cp/well), P the positive droplet number, N the negative droplet number, and V the mean volume in µL of one droplet. According to Bio-Rad, V is equal to 0.85×10^−3^ µL. Primers and probe targeted a part of the *rimM* gene, which is present in a single copy in the *Xf* genome. Therefore, the result can be directly converted into cp/µL in the initial samples, by multiplying it with the total volume of reaction mix (20 µL), and then dividing it by the volume of DNA sample added to the reaction mix (8 µL) at the beginning of the assay.

A bias, meaning the under or over-estimation of the quantification estimated by ddPCR, in comparison with the expected concentration, was calculated with the following formula:

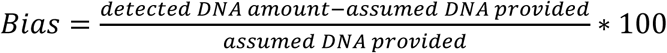

## 3 Results

### 3.1 Optimization of the ddPCR assay

The ddPCR appeared to be efficient for *Xf* detection at all tested temperatures. The first thermal gradient tested allowed us to identify 58.5°C as the most suitable hybridization temperature, for which positive droplets showed the highest fluorescence amplitude and were well distinguished from the negative droplets. The second thermal gradient confirmed this preliminary result and made it possible to fix the optimum hybridization temperature for ddPCR at 59°C. At this temperature, positive droplets presented the highest fluorescence amplitude, the less “rain” (i.e. droplets ranging between the positive and negative ones), and better separation from negative droplets (data not shown).

Addition of BSA to the reaction mix did not increase the number of droplets amplified, the number of target DNA detected, nor the amplitude of the fluorescence signal. However, as it increased the standard deviation between replicates, no BSA was added for the optimized ddPCR protocol retained in this study (data not shown).

Like for *O. europaea* shown in Figure 1, *Xf* detection was successful for all the five matrices, regardless of the volume of DNA extract added to the PCR mix (Supplemental Data 1). Increasing the DNA volume added to the mix increased the number of DNA copies detected, without complete inhibition of the reaction. As the aim of the ddPCR assay in this study was also to improve the limit of detection of *Xf* in low-level contaminated samples, the final volume of DNA chosen was 8 µL. This corresponded to the highest volume of DNA that could be added to ddPCR reaction mix in this study.

**Figure 1:**
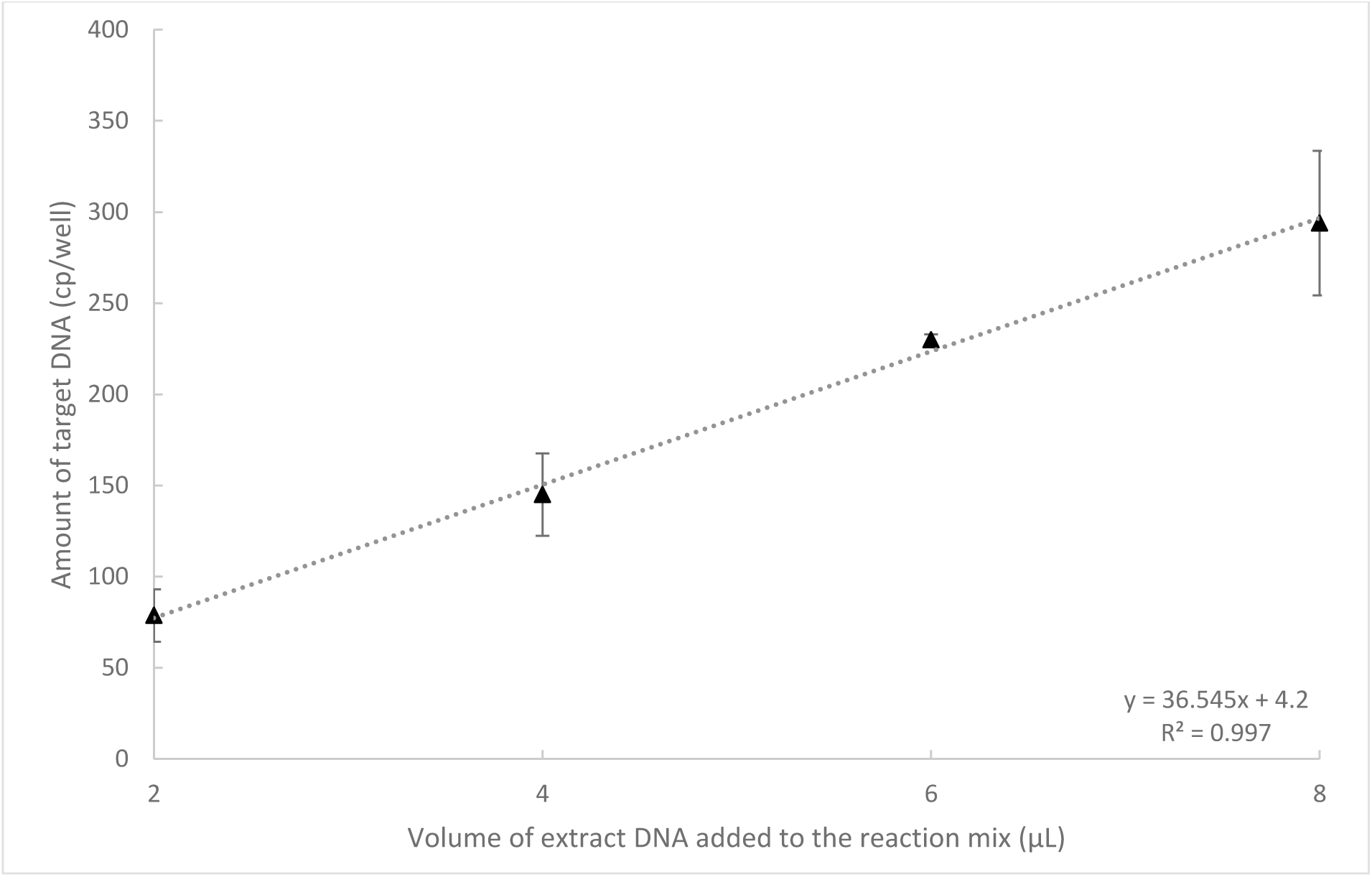
Influence of the DNA extract volume added to the ddPCR reaction mix on the amount of DNA target detected for *O. europaea*.

A tenth dilution of the DNA of *L. angustifolia* and *R. officinalis* spiked with 1×10^3^ b/mL, corresponding to the first concentration under the limit of detection obtained with ddPCR, was tested. For *L. angustifolia*, none of the six replicates provided positive droplets. In only one of the six replicates tested for *R. officinalis*, two positive droplets were detected. Dilution of the DNA extract of *L. angustifolia* and *R. officinalis* did not improve the limit of detection or was not reproducible enough in this study.

### 3.2 Xf detection by ddPCR in spiked plant samples

The healthy status of the five plants was validated before spiking, by applying real-time PCR Harper *et al*. (2010). Indeed, no Ct value was obtained for all the five plant matrices (Table 1). As expected, in these assays, no cross-reaction was found, as no positive droplets were found for any of the NTCs, for the plant matrices. With the exception of *O. europaea* (curve could not be drawn since only two concentrations gave some results), all the ddPCR matrix standard curves showed high linearity and amplification efficiency, as the correlation coefficient (R^2^) was higher than 0.96. This indicated the successful outcomes and good performances of all the assays (Table 2).

**Table 1:** Mean Ct values obtained with real-time PCR for the bacterial suspension and the five spiked plant matrices.

| Dilution range<br>(b/mL) | Bacterial suspension |  | *L. angustifolia* |  | *O. europaea* |  | *P. myrtifolia* |  | *Q. ilex* |  | *R. officinalis* |  |
| --- | --- | --- | --- | --- | --- | --- | --- | --- | --- | --- | --- | --- |
| 1x10<sup>5</sup> | 32.18 ± 0.38<sup>a</sup> | (9)<sup>b</sup> | 30.76 ± 0.18 | (9) | 36.02 ± 0.43 | (9) | 30.38 ± 0.17 | (9) | 31.85 ± 0.10 | (9) | 32.24 ± 0.17 | (9) |
| 5x10<sup>4</sup> | 33.14 ± 0.45 | (9) | 31.24 ± 0.11 | (9) | 36.59 ± 0.99 | (9) | 31.41 ± 0.18 | (9) | 32.00 ± 0.15 | (9) | 32.53 ± 0.14 | (9) |
| 1x10<sup>4</sup> | 35.46 ± 0.64 | (9) | 33.68 ± 0.20 | (9) | 37.32 ± 0.05 | (3) | 33.85 ± 0.34 | (9) | 34.90 ± 0.50 | (9) | 34.83 ± 0.66 | (9) |
| 5x10<sup>3</sup> | 35.55 ± 0.21 | (9) | 34.93 ± 0.20 | (9) | 37.86 ± 0.49 | (2) | 34.59 ± 0.28 | (9) | 35.31 ± 0.55 | (9) | 36.09 ± 1.05 | (9) |
| 1x10<sup>3</sup> | 37.42 ± 0.37 | (6) | 37.45 ± 0.58 | (9) | 37.78 ± 0.53 | (2) | 38.00 ± 0.56 | (9) | 37.62 ± 0.86 | (7) | 38.05 ± 0.50 | (6) |
| 5x10<sup>2</sup> | 38.81 ± 0.12 | (5) | 37.44 ± 0.00 | (1) | 36.61 ± 0.00 | (1) | 37.62 ± 0.18 | (5) | 37.76 ± 0.29 | (4) | 38.25 ± 0.13 | (4) |
| 1x10<sup>2</sup> | 38.00 ± 0.19 | (6) | 38.46 ± 1.01 | (2) | nd | (0) | 38.29 ± 0.13 | (4) | 36.75 ± 0.19 | (6) | nd | (0) |
| 0 | nd<sup>c</sup> | (0) | nd | (0) | nd | (0) | nd | (0) | nd | (0) | nd | (0) |

<sup>a</sup>: Average Ct ± Standard Deviation (SD)
<sup>b</sup>: Number of positive replicates on nine replicates analyzed
<sup>c</sup>: Not detected

**Table 2:** Curve information for the *Xf* bacterial suspension and the five *Xf* spiked plant matrices.

|  | Curve equation | R<sup>2</sup> |
| --- | --- | --- |
| ddPCR |  |  |
| Bacterial suspension | $y = 0.4092x + 1136.7$ | 0.99 |
| *L. angustifolia* | $y = 85x + 2\,007.35$ | 0.96 |
| *O. europaea* | N/A<sup>a</sup> |  |
| *P. myrtifolia* | $y = 0.90x - 781.02$ | 1.00 |
| *Q. ilex* | $y = 0.36x + 1\,187.91$ | 0.99 |
| *R. officinalis* | $y = 0.37x + 1\,363.74$ | 0.98 |
| REAL-time PCR |  |  |
| Bacterial suspension | $y = 2.06 \times 10^{16} e^{-8.08E-01x}$ | 0.97 |
| *L. angustifolia* | $y = 6.40 \times 10^{13} e^{-6.66E-01x}$ | 0.99 |
| *O. europaea* | $y = 1.18 \times 10^{33} e^{-1.79x}$ | 0.98 |
| *P. myrtifolia* | $y = 1.10 \times 10^{13} e^{-6.13E-01x}$ | 0.99 |
| *Q. ilex* | $y = 1.74 \times 10^{15} e^{-7.48E-01x}$ | 0.98 |
| *R. officinalis* | $y = 2.32 \times 10^{15} e^{-7.48E-01x}$ | 0.99 |

272 <sup>a</sup>: As there were results for only two concentrations ( $10^5$ b/mL and $5 \cdot 10^4$ b/mL), no curve could be drawn.

The results obtained for the five matrices and the bacterial suspension showed clear distinctions between positive (blue) and negative (grey) droplets (Figure 2). The background (negative droplets) had a similar fluorescence amplitude between samples of the same matrix, and between the five matrices (mean amplitude of fluorescence of negatives droplets ranged from 1,070 to 1,606). The threshold was manually set at an amplitude of fluorescence between 2,000 and 3,000 for each ddPCR run. Compared to the positive control, which is a lysed suspension of pure culture of *Xf*, very less rain was observed on the spiked sampled plots, showing high efficiency of the PCR reactions.

**Figure 2:**
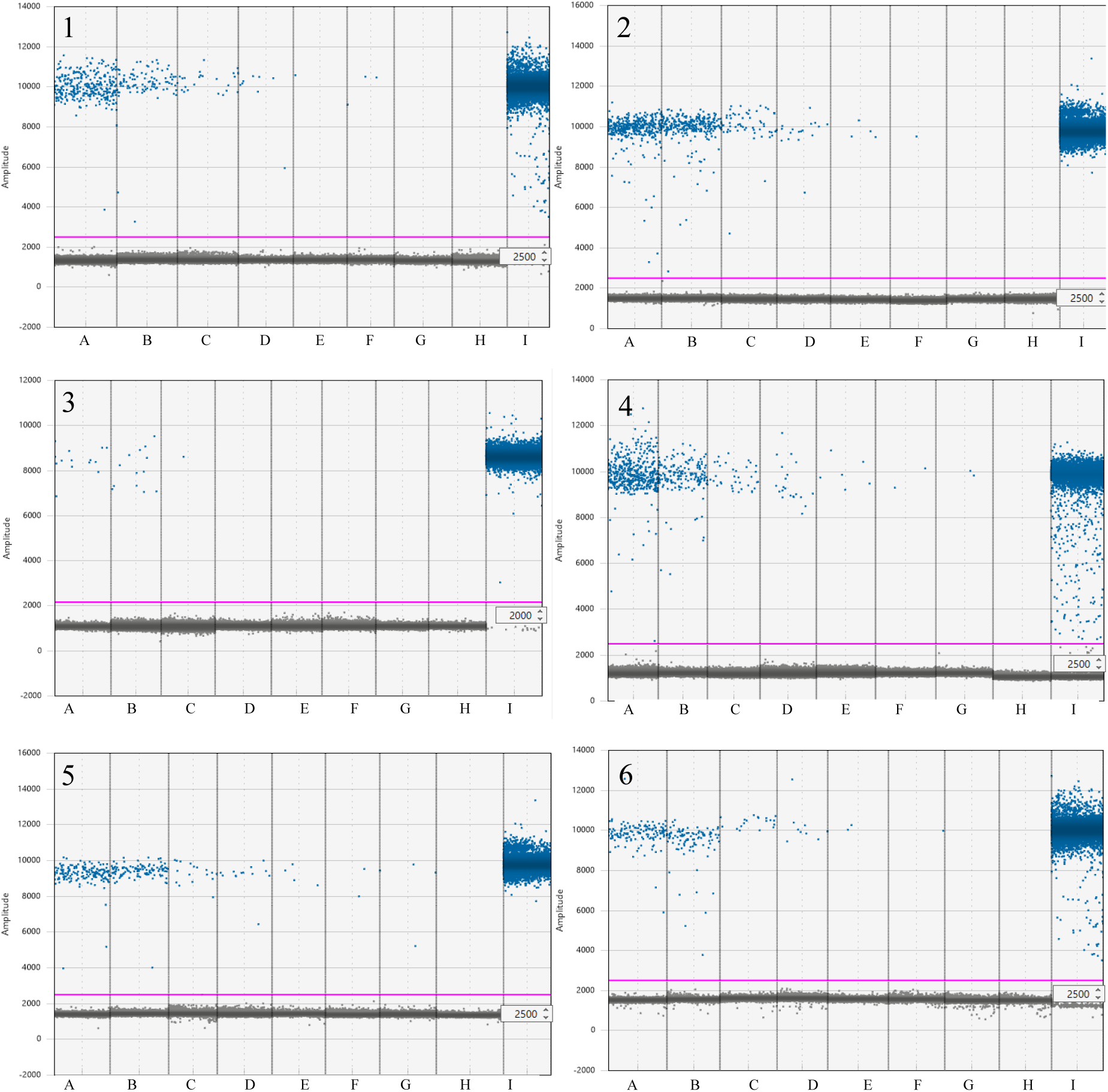
Comparison of the different limits of detection of *Xf* obtained by ddPCR in the bacterial suspension and spiked plant matrices. Pink line: threshold separating negative from positive droplets. Blue dots: positive droplets with amplification. Grey dots: negative droplets with no amplification. 1: Bacterial suspension. 2: *Lavandula* sp. 3: *O. europaea*. 4: *P. myrtifolia*. 5: *Q. ilex*. 6: *R. officinalis*. Wells A to G bacterial suspension range of *Xf*, A: 1×10^5^ b/mL; B: 5×10^4^ b/mL; C: 1×10^4^ b/mL; D: 5×10^3^ b/mL; E: 1×10^3^ b/mL; F: 5×10^2^ b/mL and G: 1×10^2^ b/mL. Well H: NTC specific to each matrix. Well I: positive control (lysis suspension of 1×10^7^ b/mL of *Xf*).

ddPCR enabled the detection of *Xf* in all the matrices, but at different concentrations (Table 3). The limit of detection was fixed at 5×10^4^ b/mL, i.e, 2.5×10^5^ b/g of plant for *O. europaea*, at 5×10^3^ b/mL, i.e, 2.5×10^4^ b/g of plant for *L. angustifolia* and *R. officinalis*, and at 1×10^3^ b/mL, i.e, 5×10^3^ b/g of plant for *Q. ilex, P. myrtifolia* and the bacterial suspension. The bias between the detected amount of DNA and the presumed amount of DNA provided was calculated for each matrix at the limit of detection. Compared to the quantity of DNA expected, DNA quantifications of *Xf* were overestimated by 6.54% and 23.96% in *Q. ilex* and in the bacterial suspension, respectively. In *L. angustifolia, R. officinalis, P. myrtifolia*, and *O. europaea*, the DNA quantifications of *Xf* were underestimated by 3.08%, 24.36%, 32.03% and 95.60%, respectively.

**Table 3:** Mean concentrations estimated in copies/mL (cp/mL) obtained with ddPCR for the bacterial suspension and the five spiked plant matrices.

| Dilution range<br>(b/mL) | Bacterial suspension |  | *L. angustifolia* |  | *O. europaea* |  | *P. myrtifolia* |  | *Q. ilex* |  | *R. officinalis* |  |
| --- | --- | --- | --- | --- | --- | --- | --- | --- | --- | --- | --- | --- |
| 1x10<sup>5</sup> | 4.02x10<sup>4</sup><sup>a</sup><br>± 3.63x10<sup>3</sup> | (3/3)<sup>b</sup> | 8.07x10<sup>4</sup><br>± 3.70x10<sup>3</sup> | (3/3) | 3.38x10<sup>3</sup><br>± 5.67x10<sup>2</sup> | (9/9) | 9.08x10<sup>4</sup><br>± 4.95x10<sup>3</sup> | (3/3) | 3.58x10<sup>4</sup><br>± 2.96x10<sup>3</sup> | (3/3) | 3.62x10<sup>4</sup><br>± 3.71x10<sup>3</sup> | (3/3) |
| 5x10<sup>4</sup> | 2.53x10<sup>4</sup><br>± 2.20x10<sup>3</sup> | (3/3) | 5.81x10<sup>4</sup><br>± 2.02x10<sup>3</sup> | (3/3) | 2.20x10<sup>3</sup><br>± 5.45x10<sup>2</sup> | (9/9) | 4.03x10<sup>4</sup><br>± 4.71x10<sup>3</sup> | (3/3) | 2.30x10<sup>4</sup><br>± 9.92x10<sup>2</sup> | (3/3) | 2.38x10<sup>4</sup><br>± 3.20x10<sup>3</sup> | (3/3) |
| 1x10<sup>4</sup> | 5.73x10<sup>3</sup><br>± 1.14x10<sup>3</sup> | (3/3) | 1.17x10<sup>4</sup><br>± 2.19x10<sup>3</sup> | (3/3) | 5.28x10<sup>2</sup><br>± 1.90x10<sup>2</sup> | (4/9) | 6.17x10<sup>3</sup><br>± 1.39x10<sup>3</sup> | (3/3) | 4.95x10<sup>3</sup><br>± 4.04x10<sup>2</sup> | (3/3) | 5.49x10<sup>3</sup><br>± 2.98x10<sup>2</sup> | (3/3) |
| 5x10<sup>3</sup> | 2.84x10<sup>3</sup><br>± 8.78x10<sup>2</sup> | (9/9) | 4.85x10<sup>3</sup><br>± 1.21x10<sup>3</sup> | (3/3) | 4.25x10<sup>2</sup><br>± 0.00 | (1/9) | 5.14x10<sup>3</sup><br>± 1.48x10<sup>3</sup> | (3/3) | 2.81x10<sup>3</sup><br>± 7.90x10<sup>2</sup> | (9/9) | 3.78x10<sup>3</sup><br>± 2.16x10<sup>3</sup> | (9/9) |
| 1x10<sup>3</sup> | 8.54x10<sup>2</sup><br>± 2.57x10<sup>2</sup> | (9/9) | 5.66x10<sup>2</sup><br>± 1.63x10<sup>2</sup> | (6/9) | No Call | (0/9) | 6.80x10<sup>2</sup><br>± 2.68x10<sup>2</sup> | (9/9) | 1.07x10<sup>3</sup><br>± 5.58x10<sup>2</sup> | (9/9) | 9.04x10<sup>2</sup><br>± 1.61x10<sup>2</sup> | (7/9) |
| 5x10<sup>2</sup> | 6.20x10<sup>2</sup><br>± 1.19x10<sup>2</sup> | (8/9) | No Call | (0/9) | No Call | (0/9) | 5.15x10<sup>2</sup><br>± 1.43x10<sup>2</sup> | (4/9) | 7.17x10<sup>2</sup><br>± 2.59x10<sup>2</sup> | (7/9) | 6.18x10<sup>2</sup><br>± 4.56x10<sup>2</sup> | (5/9) |
| 1x10<sup>2</sup> | 5.84x10<sup>2</sup><br>± 2.06x10<sup>2</sup> | (6/9) | 3.73x10<sup>2</sup><br>± 0.00 | (1/9) | No Call | (0/9) | 1.77x10<sup>2</sup><br>± 0.00 | (1/9) | 6.23x10<sup>2</sup><br>± 1.50x10<sup>2</sup> | (7/9) | 2.01x10<sup>2</sup><br>± 2.92x10<sup>1</sup> | (2/9) |
| 0 | No Call | (0/9) | No Call | (0/9) | No Call | (0/9) | No Call | (0/9) | No Call | (0/9) | No Call | (0/9) |

<sup>a</sup>: Average Ct ± SD
<sup>b</sup>: Number of positive replicates/number of replicates analyzed

### 3.3 Real-time PCR vs ddPCR for Xf detection in spiked samples

As for ddPCR, all the real-time PCR matrix standard curves showed high linearity and amplification efficiency, as the correlation coefficient (R^2^) was greater than 0.97. This indicated the successful outcomes and good performances of all the assays (Table 2).

Real-time PCR Harper *et al*. (2010) was used as a reference method in this study. For all the assays, no mean Ct values exceeding 38 were recorded at a concentration equal to or higher than the limit of detection, meaning that the limit of detection and positive results were consistent. Moreover, for *O. europaea* the EPPO PM 7/24 protocol mentions a limit of detection for samples artificially contaminated with *Xf* subsp. *multiplex* of 100% at 1×10^5^ b/mL. In this study, the limit of detection of 5×10^4^ b/mL was close to that presented in the PM 7/24 (EPPO, 2018b). The limit of detection of *Xf* in *P. myrtifolia* is known to be 1×10^3^, which is the same as the value we found (Legendre B., personal communication). The limits of detection of *Xf* subsp. *multiplex* for the other matrices could not be compared, as there are no available data.

Real-time PCR and ddPCR technology provided equivalent limits of detection for *Xf* in the following matrices: *O. europaea, P. myrtifolia* and *R. officinalis* (Table 1, Table 3). The ddPCR technology presented a slightly higher limit of detection of 0.5 log for *L. angustifolia*. However, a decrease in the limit of detection for *Xf* of 0.5 log for *Q. ilex* and the bacterial suspension wereobserved. In the conditions of DNA extraction used for this study, and according to the volume of DNA added to the real-time PCR assay, the theoretical limit of detection should be 1×10^2^ b/mL for the five plant matrices. These results revealed that *L. angustifolia* and *P. myrtifolia* may contain fewer real-time PCR inhibitors than *Q. ilex* and *R. officinalis*. Moreover, the limit of detection of the bacterial suspension was 5×10^3^ b/mL, meaning that the QuickPick extraction kit may not be 100% efficient to extract the DNA of bacteria in pure culture.

### 3.4 Xf detection in naturally infected samples: real-time PCR vs ddPCR

A total of 22 samples from infected areas were tested using real-time PCR and ddPCR. Of these, 20 had a mean Ct value below 38 (ranging from 23.30 to 37.00) (Table 4). However, two samples, P13 and Q04, had a Ct value above 38 (38.65 and 39, respectively), and were considered negative.

**Table 4:**
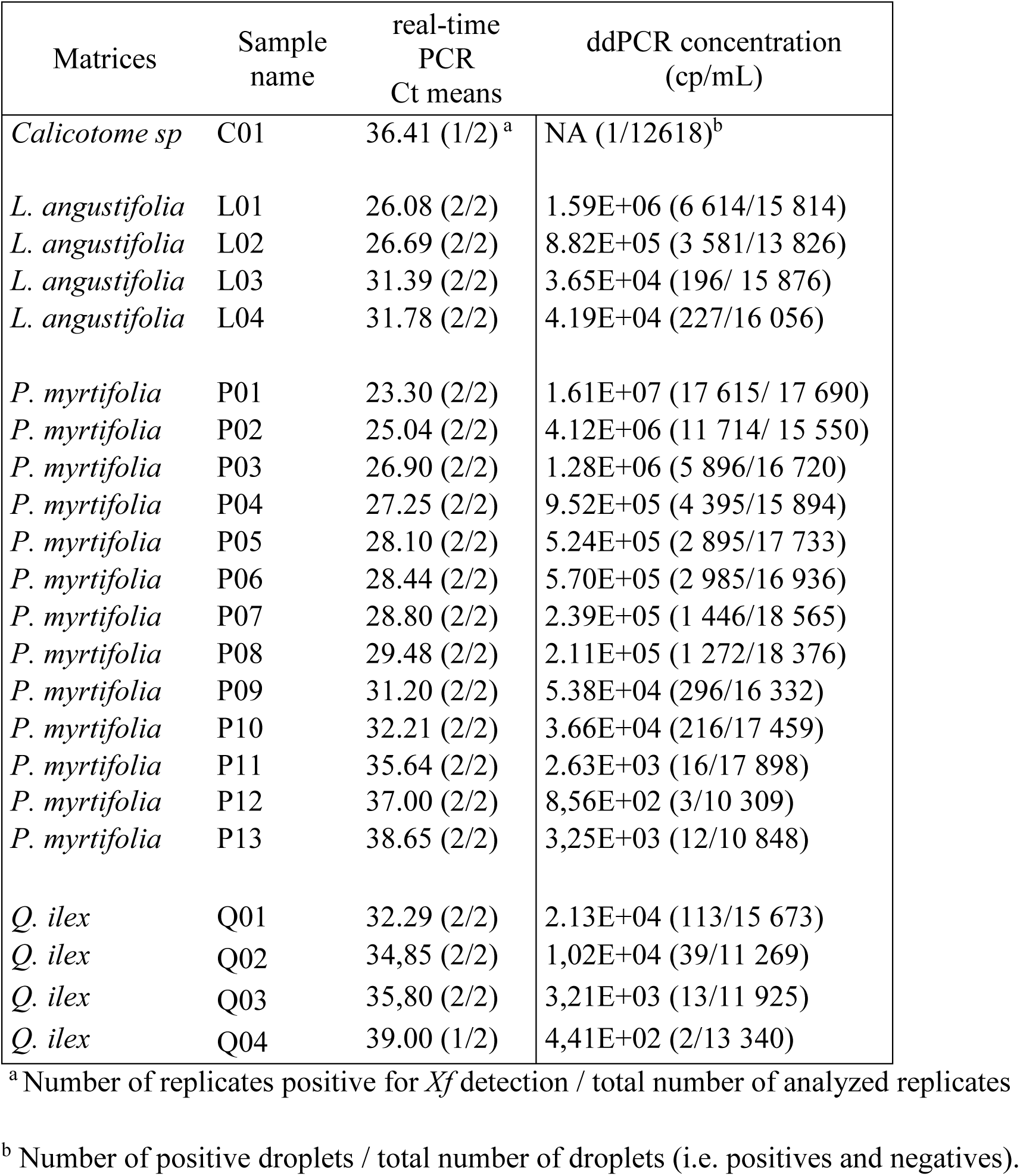
Mean Ct values and concentrations estimated in cp/mL of naturally infected samples obtained after real-time PCR and ddPCR.

The 22 naturally infected samples were then analyzed by ddPCR. The presence of *Xf* was detected in 21 of them, including samples P13 and Q04, with at least two droplets, and a total concentration ranging from 4.41×10^2^ cp/mL to 1.61×10^7^ cp/mL, confirming the ability of ddPCR to detect *Xf* in naturally infected samples. As only one positive droplet was detected for sample C01, this sample was considered negative by ddPCR. With the exception of samples L04, P06 and P13, the decrease in the Ct value was correlated with an increase in the quantity of DNA detected by ddPCR (Table 4). For each matrix the results obtained by ddPCR and real-time PCR were compared and were highly correlated (Figure 3).

**Figure 3:**
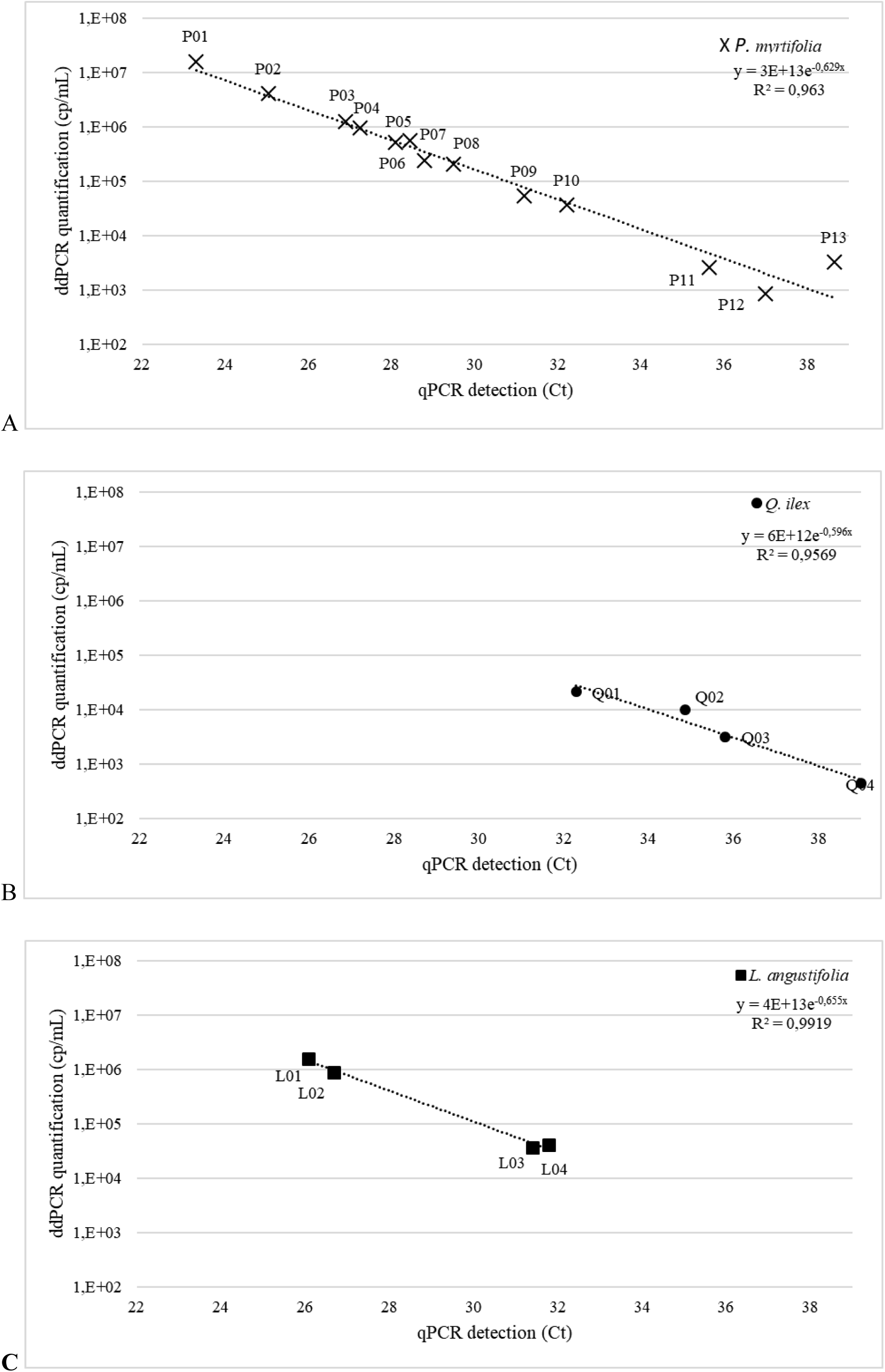
Correlation between the Ct values obtained by real-time PCR and the amount of target DNA quantified by ddPCR (log cp/mL) for the naturally infected samples analyzed. A: samples of *P. myrtifolia*; B: samples of *Q. ilex*; C: samples of *L. angustifolia*.

## 4 Discussion

It has been observed that tenfold dilutions of the extracted DNA could reduce the effects of real-time PCR inhibitors (Francis *et al*., 2006; Minsavage *et al*., 1994; Modesti *et al*., 2017). In this study, diluting the DNA extract of *L. angustifolia* and *R. officinalis* did not reduce the limit of detection using ddPCR, or the obtained results were not sufficiently reproducible. This approach does not seem to be useful and appropriate for *Xf* detection in these two matrices, using ddPCR. Nevertheless, more tests should be carried out to support this assumption.

Compared to real-time PCR, ddPCR can be considered as a controversial method. Some studies have revealed that ddPCR was useful to improve pathogen detection sensitivity and to decrease the impact of PCR inhibitors on PCR efficiency (Arvia *et al*., 2017; Bharuthram *et al*., 2014; Dong *et al*., 2018; Rački *et al*., 2014; Zhao *et al*., 2016). In other cases, ddPCR was 10 and 100 fold less sensitive than real-time PCR in detecting cytomegalovirus and *Leishmaniasis* parasite DNA, respectively (Hayden *et al*., 2013; Ramírez *et al*., 2019). Dreo *et al*., reported that ddPCR benefits were dependent of the pathosystem studied (Dreo *et al*., 2014). The detection of *Erwinia amylovora* showed similar levels using real-time PCR and ddPCR, while the detection of *R. solanacearum* in low-level infected samples was improved by ddPCR (Dreo *et al*., 2014). In our study, the two methods showed the same limit of detection for *O. europaea, P. myrtifolia* and *R. officinalis*. Real-time PCR allowed better detection of 0.5 log for *L. angustifolia*, and ddPCR allowed better detection of 0.5 log for *Q. ilex* and bacterial suspension.

ddPCR was also compared with real-time PCR on 22 naturally infected samples. Using real-time PCR, two samples P13 and Q04 had a Ct value higher than 38, and were thus considered negative, i.e. not infected by *Xf*. As these two samples were frozen at −20°C for one year, it is possible that the DNA was altered, explaining the higher Ct values obtained in this study. The other 20 samples, considered positive, had a Ct value lower than 38. Using ddPCR, *Xf* was detected in 21 samples, as more than two positive droplets were obtained. ddPCR did not reveal the presence of *Xf* in sample C01 of *Calicotome* sp., unlike real-time PCR that detected *Xf* in only one of the duplicates tested. These results could highlight the presence of PCR inhibitors in this matrix. Moreover, ddPCR technology enabled the detection of *Xf* in both samples P13 and Q04, considered in this study as not infected by *Xf* using real-time PCR. As shown by Dreo *et al*. (2014) for the detection of *R. solanacearum*, ddPCR technology could offer a real advantage for the detection of pathogenic bacteria, and can be applied to the detection of *Xf* in contaminated plants with low concentrations of target DNA (Dreo *et al*., 2014). It also successfully confirmed the positive samples identified using real-time PCR.

## 5 Conclusion

In this work, we proposed the first suitable ddPCR assay for the detection of *Xf* in plants. We easily transferred a well-known routinely used real-time PCR technique for *Xf* detection in ddPCR. Here, we reported all the set up steps leading to the optimal protocol and its comparison with the current routine method. The results demonstrated the usefulness of ddPCR technology as an alternative method for *Xf* detection in plants. However, the ddPCR assay is more time-consuming than real-time PCR and does not seem to be suitable for routine analysis. This technology requires more steps than real-time PCR. Furthermore, the reaction mix has to handle with care to ensure the generation of the appropriate number of generated droplets. Nevertheless, as only two droplets are needed to confirm a sample as positive with ddPCR, this method could confirm the status of samples found to be negative by real-time PCR due to high Ct values, and could improve *Xf* detection in low-level infected samples. ddPCR should be tested on insects to see whether this technology would still be efficient, and whether it offers a benefit for *Xf* detection in this matrix.

## 6 Acknowledgements

We thank Marie-Agnès Jacques, Philippe Reignault, Pascal Gentit and Mathieu Rolland for fruitful discussions and critical reading of the manuscript.

