## Supplemental data 1 for "Comparison of real-time PCR and droplet digital PCR for the detection of *Xylella fastidiosa* in plants"

|  |  | **Amount of DNA target detected (cp/well)** | | | | | |
| --- | --- | --- | --- | --- | --- | --- | --- |
| **DNA volume**  **(µL)** |  | **Bacterial**  **suspension** | ***L. angustifolia*** | ***O. europaea*** | ***P. myrtifolia*** | ***Q. ilex*** | ***R. officinalis*** |
| 2 |  | 109,0 | 104,0 | 78,7 | 57,1 | 80,9 | 89,9 |
| 4 |  | 164,5 | 257,0 | 145,0 | 123,0 | 143,5 | 134,5 |
| 6 |  | 276,0 | 342,0 | 230,0 | 190,0 | 219,5 | 222,5 |
| 8 |  | 301,5 | 443,5 | 294,0 | 229,7 | 261,5 | 272,0 |

**Supplemental data 1: Influence of the DNA extract volume added to the ddPCR reaction mix on the amount of DNA target detected.**
